# Neutral genetic diversity in mixed mating systems

**DOI:** 10.1101/2022.07.29.502020

**Authors:** Marcy K. Uyenoyama

**Affiliations:** Department of Biology, Box 90338, Duke University, Durham, NC 27708-0338, USA

**Author notes:** Address for correspondence: Marcy K. Uyenoyama, Department of Biology, Box 90338, Duke University, Durham, NC 27708-0338, USA.

**Keywords:** effective number, Ewens Sampling Formula, selfing, hermaphroditism, mating system, conservation genetics

## Abstract

Systems of reproduction differ with respect to the magnitude of neutral genetic diversity maintained in a population. In particular, the partitioning of reproductives into mating types and regular inbreeding have long been recognized as key factors that influence effective population number. Here, a range of reproductive systems (full gonochorism, full hermaphroditism, androdioecy, and gynodioecy) are compared with respect to the maintenance of neutral genetic diversity. The analysis assumes anisogamy, with reproduction limited by the availability of large gametes (ova or seeds) but not small gametes (sperm or pollen). Levels of neutral genetic diversity respond to the relative proportions of gonochores and hermaphrodites in different ways under androdioecy versus gynodioecy. The manner in which effective number, sex-specific viability differences, and the evolving quantitative trait of the population influence the level of neutral genetic diversity is described across the systems of reproduction studied.

## 1 Introduction

Genomic patterns of neutral variation reflect the evolutionary context that generated them, including selection targeted to linked and unlinked genomic locations (Stephan 2019), population structure (Barton 2008), and the system of reproduction. This analysis addresses various aspects of life history that can influence levels of neutral diversity across the genome, including inbreeding, inbreeding depression, subdivision of reproductives into mating types (sexes), the evolution of the sex ratio, and sex-specific viability.

### 1.1 Factors influencing the level of diversity

Inbreeding tends to reduce diversity by promoting mating between relatives bearing similar genes. Inbreeding depression and other processes may further reduce diversity by decreasing the number of reproductive individuals. The models considered here incorporate inbreeding through perhaps the simplest mechanism of self-fertilization (selfing) by hermaphrodites. One might expect that forms of biparental inbreeding in gonochorous organisms may induce qualitatively similar effects on the level of neutral diversity.

Under systems of reproduction in which only partners of different mating types can generate offspring, evolutionary pressures on the sex ratio (relative numbers of mating types) may also affect the maintenance of neutral diversity. An evolutionarily stable strategy (ESS) corresponds to an “unbeatable” population sex ratio (relative numbers of mating types) (Hamilton 1967), from which no modification can increase at a geometric rate (see Maynard Smith 1978, Chapter 9). Even if the population sex ratio at conception achieves the ESS, sex-specific viability may skew the sex ratio at reproductive age and the relative contributions of the mating types to the next generation.

To assess the effects of these major determinants of the level of neutral diversity maintained in populations, this study presents a coalescence-based analysis of a range of reproductive systems, including full gonochorism, full hermaphroditism, androdioecy, and gynodioecy. A single index, relative effective number (Section 2.1) summarizes the effects of these factors on the maintenance of neutral diversity.

### 1.2 Systems of reproduction

All reproductive systems studied here are anisogamous, with zygotes formed by fusion between a female gamete (large) and a male gamete (small). Full gonochorism implies mating exclusively between distinct mating types, each of which specializes in the production of a single gamete type.

Hermaphroditism introduces the possibility of self-fertilization: the production of offspring through fusion of gametes generated by a single individual. In the models considered here, outcrossing rate 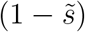 denotes the proportion of large gametes that are fertilized by a male gamete randomly derived from the population.

#### Androdioecy

Androdioecious populations comprise 2 mating types: males, which produce only small gametes, and hermaphrodites, which produce both large and small gametes (Figure 1). Model systems include the plant *Datisca glomerata* (Wolf *et al*. 2001) and the nematode *Caenorhabditis elegans* (Steward and Phillips 2002). Androdioecy is rare among plants as well as animals. In the models considered here, each male contributes to the population pool of male gametes (denoted as pollen) at rate *σ*_*A*_ relative to each hermaphrodite.

**Figure 1:**
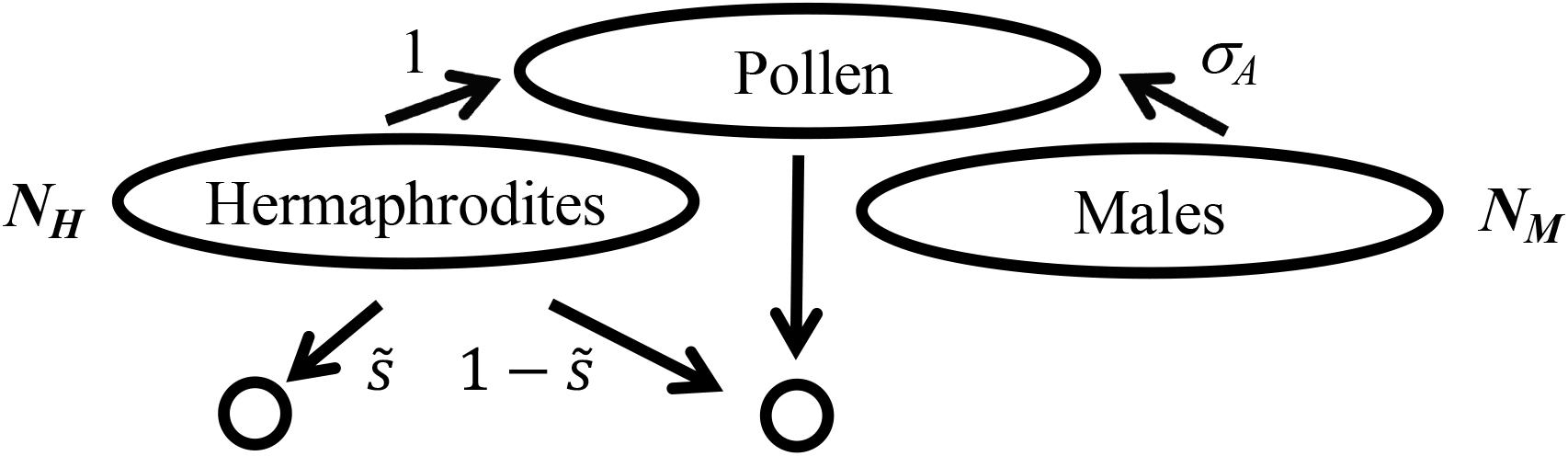
Offspring production under androdioecy. All female gametes derive from hermaphrodites. A proportion 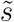 of newly-formed zygotes derive from the fusion between a female gamete and a male gamete produced by the same hermaphrodite, and 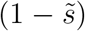 derive from pollen sampled from the population. Each of the *N*_*M*_ reproductive males contributes pollen at rate *σ*_*A*_ relative to each of the *N*_*H*_ reproductive hermaphrodites.

#### Gynodioecy

Gynodioecious taxa, comprising hermaphrodites together with females, appear to derive from multiple independent origins in flowering plants (Rivkin *et al*. 2016). Only hermaphrodites produce small gametes, which fertilize large gametes produced by both females and hermaphrodites. In the models considered here, a female contributes *σ*_*G*_ female gametes per female gamete contributed by a hermaphrodite (Figure 2). The expression of any positive level of male fertility by hermaphrodites or males is assumed to be fully sufficient to maintain the population. This assumption gives rise to quantitative and qualitative differences in the level of neutral genetic diversity maintained by the reproductive systems studied here.

**Figure 2:**
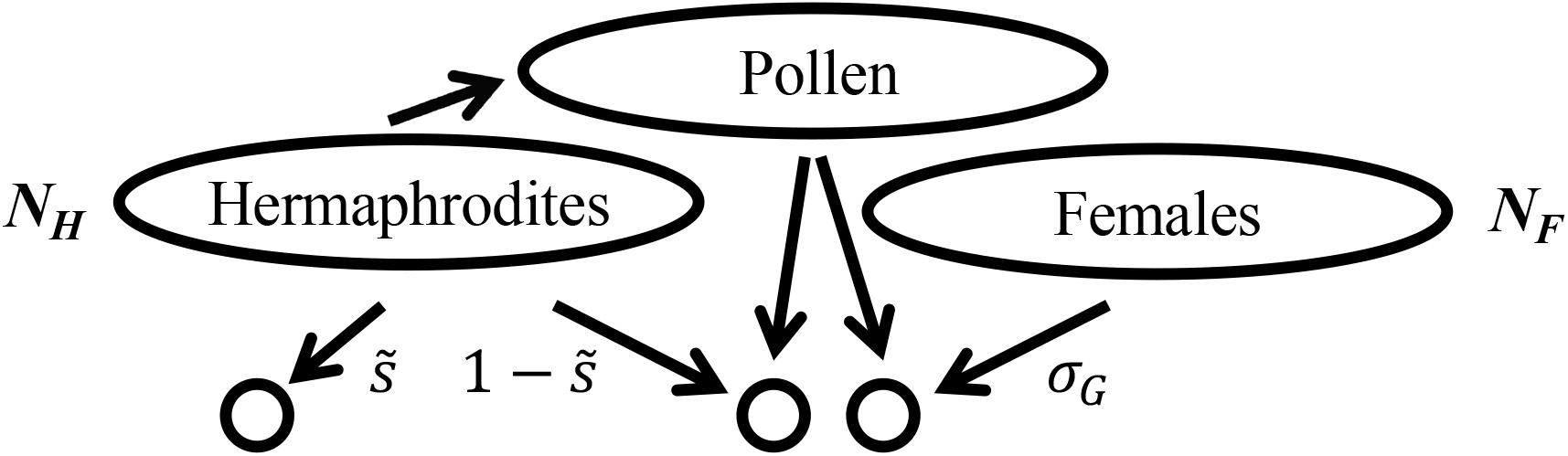
Offspring production under gynodioecy. Hermaphrodites alone contribute to the population pollen pool. As for androdioecy (Fig. 1), hermaphrodites set a proportion 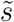 of their seeds (female gametes) exclusively by self-pollen and the complement by the population pollen pool. Relative to a hermaphrodite, each of the *N*_*F*_ females generates *σ*_*G*_ zygotes through seeds, all which are fertilized from the population pollen pool.

## 2 Key components of the evolutionary process

This section presents an overview of the analytical approaches used in assessing the effects of inbreeding, reproductive value of mating types, evolution of the sex ratio, and differential viability on relative effective number *R* (2). These concepts are illustrated here through the familiar cases of full hermaphroditism and full gonochorism. They form the basis of the analysis of dioecious mating systems, in which the several components interact (Sections 3.1 and 3.2).

### 2.1 Relative effective number

Relative effective number (*R*), the ratio of two kinds of effective number, serves as the basis for comparison across reproductive systems with respect to the maintenance of neutral genetic diversity.

Sewall Wright introduced many definitions of effective number, denoting a variety of factors influencing genetic diversity (*e*.*g*., Wright 1939). That some have divergent expressions within a given model (Ewens 1982; Caballero and Hill 1992) reflects that the various definitions address distinct aspects of the evolutionary process.

Let 1*/N*_*P*_ denote the rate of parent-sharing, the probability that a pair of genes randomly sampled from distinct individuals derive from the same parent. In the substantial literature on effective number, *N*_*P*_ has been called “inbreeding effective number” (Crow 1954; Crow and Denniston 1988; Ewens 1982). In addition, coalescence effective number is denoted by *N*_*C*_, with 1*/*2*N*_*C*_ the per-generation rate at which a pair of lineages randomly sampled from distinct individuals coalesce in their most recent common ancestor.

Under the usual large-population approximation of coalescence theory, the scaled mutation rate corresponds to

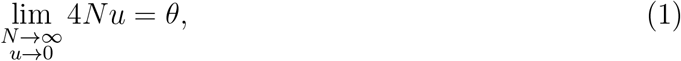

in which *u* is the per-generation rate of neutral mutation. Beyond the requirement that *N* and 1*/u* go to infinite at the same rate, scaling factor *N* is largely arbitrary. A simple approach assigns *N* as the total number of reproductives that contribute to the next generation. For example, *N* corresponds to (*N*_*H*_ + *N*_*M*_) under androdioecy (Figure 1) and to (*N*_*H*_ + *N*_*F*_) under gynodioecy (Figure 2).

This analysis uses relative effective number (*R*) as a standard of comparison among systems of reproduction, with the total number of reproductives (*N*) and the scaled rate of mutation (*θ*) held constant. Relative effective number corresponds to the ratio of *N*_*C*_ (determined from the rate of coalescence) to *N* :

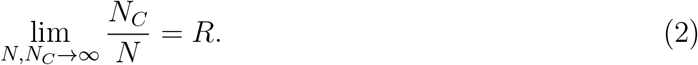

Motivation for this index of diversity derives from the Ewens Sampling Formula (ESF, Ewens 1972), which provides a succinct summary under the infinite-alleles model of mutation of the pattern of neutral genetic diversity observed in a sample of arbitrary size. The state of a sample comprising *n* genes corresponds to an allele frequency spectrum (AFS) of the form

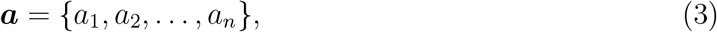

in which *a*_*i*_ denotes the number of allelic classes that appear with multiplicity *i* in the sample. Sample size is given by

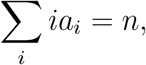

and the number of distinct alleles in the sample by

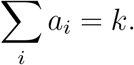

The allele frequency spectrum (AFS) differs from the site frequency spectrum (SFS), which refers to the number of observations of an allele in a biallelic sample. While the SFS distribution can be derived in various ways (*e*.*g*., Fu 1995), a simple and revealing approach involves conditioning the ESF on the observation of *K* = 2 alleles in the sample (Ganapathy and Uyenoyama 2009).

While Ewens’s (1972) original paper addressed a panmictic population, Griffiths and Lessard (2005) showed that certain models incorporating population structure converge to the Kingman (1982) coalescent process under an appropriate rescaling of time. Relative effective number (*R*) corresponds to the key scaling factor in the sense that the forms of population structure studied here can be accommodated by multiplying the mutation parameter (*θ*) in the classical ESF by *R*. The ESF, with *Rθ* substituted for *θ*, provides the probability of observing AFS ***a*** in a random sample of *n* genes:

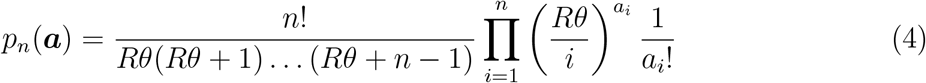

(Appendix A), with the set of parameters governing evolutionary change corresponding to

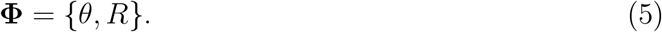

For a given value of the scaled mutation rate *θ*, higher *R* implies higher diversity: greater numbers of alleles observed in samples of a given size and an increased probability that the next-sampled gene represents an allelic class not yet observed in the sample.

### 2.2 Relative effective number under full gonochorism

To illustrate the determination of the key index of diversity *R* (2), I address the case of full gonochorism with two sexes, female and male.

Parent-sharing between a random pair of autosomal genes sampled from distinct offspring of reproductive age requires that either both genes be of maternal origin or both of paternal origin. Let *P* denote the collective contribution of the *N*_*F*_ female reproductives to the set of genes transmitted to offspring and (1 − *P*) the collective contribution of the *N*_*M*_ male reproductives. The probability that a pair of genes randomly sampled from distinct offspring of reproductive age corresponds to

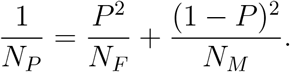

For autosomal genes, offspring receive half their genome from each parent (*P* = 1*/*2), implying

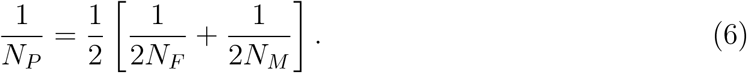

This expression indicates that the effective number of reproductives corresponds to the harmonic mean of twice the number of females and twice the number of males. This result has been derived under various definitions of effective number (*e*.*g*., Wright 1931; Ewens 1982; Caballero 1994). Crow and Kimura (1970, p. 102) appear to have presented the earliest retrospective (coalescent) argument that refers explicitly to the probability of parent-sharing.

Descent of gene pair from the same gene in the parental generation also entails that both genes be of maternal origin or of paternal origin:

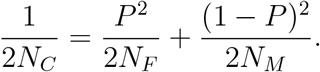

Because the effective numbers implied by the rates of parent-sharing and coalescence are in fact equal (*N*_*P*_ = *N*_*C*_), relative effective number (2) corresponds to

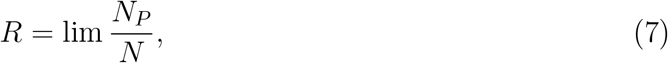

for *N* the total number of reproductive individuals

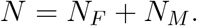

Relative effective number *R* clearly depends on the sex ratio among reproductives. Fisher (1930) addressed the evolution of the sex ratio in the context of reproductive value. Each reproducing female has a reproductive value of

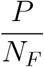

and each reproducing male,

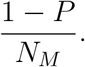

An evolutionarily stable strategy (ESS) corresponds to an “unbeatable” trait value (Hamilton 1967), at which no alleles that modify the trait value increase at geometric rates (see Maynard Smith 1978, Chapter 9). A heuristic marginal argument (Fisher 1958, p. 158) suggests that the ESS sex ratio corresponds to the point at which the returns on investing a unit of reproduction in each sex are equal:

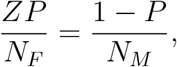

in which *Z* denotes the number of female reproductives that can be generated per male reproductive. This argument implies an ESS sex ratio among reproductives of

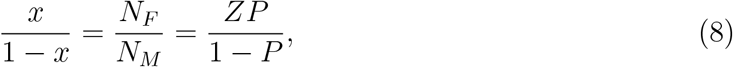

for *x* the proportion of females. For each of the mixed mating models addressed here, the ESS implied by this argument does in fact agree with the unbeatable sex ratio determined from full dynamic analyses of the evolution of autosomal sex ratio modifiers (Uyenoyama and Takebayashi 2017). For other genomic locations of the sex ratio modifier (especially sex-linkage), the ESS reflects asymmetries in relatedness of the modifier to each sex (Uyenoyama and Feldman 1981).

In general, *Z* reflects various forms of sex-specific demands by offspring on parental resources. For simplicity, I restrict consideration here to *Z* deriving solely from sex-specific viability: *Z* is the rate at which females survive from conception to reproductive age relative to males. In this case, Fisher (1958) noted that the ESS sex ratio at conception corresponds to

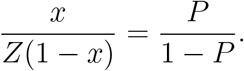

From the perspective of autosomal modifiers expressed in parents, parents are equally related to offspring of the two sexes (*P* = 1*/*2). The ESS sex ratio at conception is then unity, irrespective of the value of *Z*.

The effective numbers that determine *R* (7) (including *N*_*H*_, *N*_*M*_, and *N*_*F*_ in Figures 1 and 2) correspond to numbers of individuals at reproductive age rather than at conception. Accordingly, differential viability among mating types (*Z* ≠ 1) influences the level of neutral diversity maintained in populations. Substitution of the relative frequencies of female (*x*) and male (1 − *x*) reproductives at the ESS (8) into (6) produces

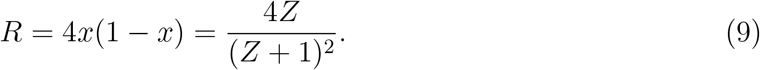

Relative effective number achieves its maximum value (*R* = 1) only in the absence of sex-specific differences in viability (*Z* = 1). For a given scaled mutation parameter (*θ*) and total number of reproductives (*N*), disparity between the sex ratio at conception and the sex ratio at reproductive age uniformly reduces the level of neutral genetic diversity maintained.

### 2.3 Effect of selfing

Reproduction by hermaphrodites introduces the possibility of self-fertilization (selfing) and the expression of inbreeding depression.

Under full hermaphroditism, as for androdioecy and gynodioecy (Figs. 1 and 2), 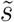 denotes the fraction of seeds that each of *N* reproductive hermaphrodite sets immediately by self-pollen, with the complement 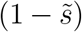 set by pollen sampled from the local gamete pool. The proportion of newly-formed zygotes that are uniparental corresponds to

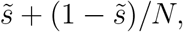

reflecting the generation of uniparental zygotes among the seeds set by pollen sampled from the population at large. Let *s*_*u*_ denote the *uniparental fraction*, the probability that a random individual of reproductive age (rather than at conception) is uniparental. Here, I assume that reproduction occurs subsequent to phases in the life cycle in which inbreeding depression or other forms of differential viability are expressed. Under full hermaphroditism, the uniparental fraction corresponds to

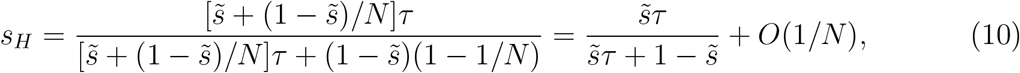

for *τ* the rate of survival of uniparental offspring relative to biparental offspring. The uniparental fraction corresponds to *s*_*A*_ (24) under androdioecy, and to *s*_*G*_ (33) under gynodioecy.

Derivation of relative effective number *R* (2) entails determining the per-generation rates of coalescence (1*/*2*N*_*C*_) and parent-sharing (1*/N*_*P*_). As coalescence entails descent of a gene pair both from the same parent and from the same gene in that parent, these rates are clearly closely related. Under androdioecy and gynodioecy as well as full hermaphroditism, the rate of coalescence corresponds to

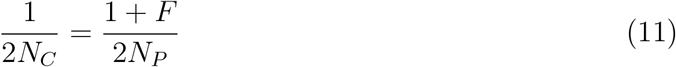

(see Uyenoyama 2024), for *F* Wright’s correlation between uniting gametes:

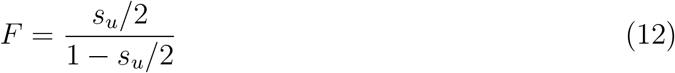

(Pollak 1987). Relative effective number (2) corresponds to

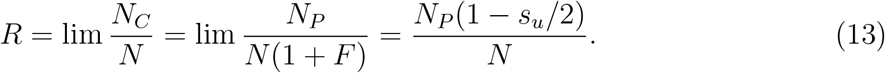

Equality between the rates of coalescence and parent-sharing (*N*_*C*_ = *N*_*P*_) holds in the absence of inbreeding (*F* = 0).

Under full hermaphroditism, a pair of genes randomly sampled from distinct reproductive individuals derive from the same parent with probability 1*/N*, irrespective of the uniparental fraction:

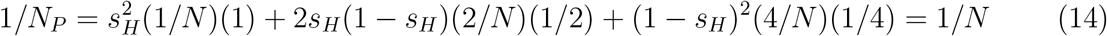

(Appendix B). In this case (*N*_*P*_ = *N*), (13) implies that

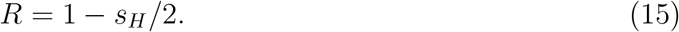

In dioecious reproductive systems, however, the rate of parent-sharing 1*/N*_*P*_ itself generally depends on the uniparental fraction *s*_*u*_.

### 2.4 Relative effective number in dioecious systems

I now address relative effective number *R* (13) in dioecious systems in which *N*_*H*_ hermaphrodites and *N*_*G*_ gonochores (females or males) contribute to the next generation. Let *P* now denote the proportion of the gene pool contributed by gonochores and (1 − *P*) the proportion contributed by hermaphrodites. As under full gonochorism (Section 2.2), the probability of parent-sharing between a random pair of genes sampled from distinct reproductive individuals depends on the proportion of the gene pool that derives from each mating type:

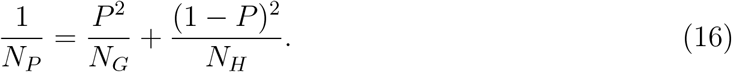

Because gonochores necessarily outcross to hermaphrodites,

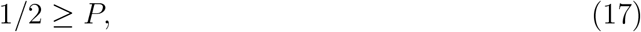

with equality only for cases in which hermaphrodites are also obligated to outcross to gonochores.

Under the heuristic reproductive value argument presented for full gonochorism, the ESS sex ratio corresponds to the point of equality between the marginal reproductive values of gonochores and hermaphrodites:

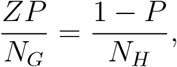

for *Z* the number of gonochoristic reproductives that can be generated per hermaphroditic reproductive. The ESS ratio of gonochores to hermaphrodites implied by this argument is

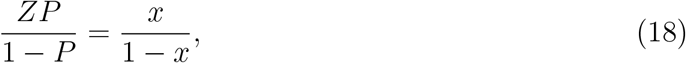

for *x* the proportion of gonochores among reproductives.

At the ESS sex ratio (18), relative effective number (13) corresponds to

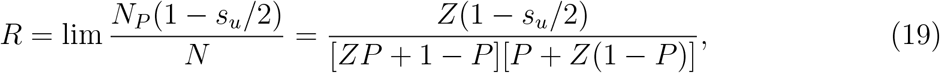

for *s*_*u*_ the probability that a random reproductive individual is uniparental This expression indicates that

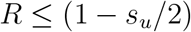

(compare (15)), with equality only at *P* = 0 (full hermaphroditism) or *Z* = 1 (absence of sex-specific viability).

While relative effective number *R* depends on the uniparental fraction alone (15) under full hermaphroditism, it may depend on all model parameters through their effects on the collective genetic contribution of gonochores (*P*) in dioecious systems. Under the chain rule, the derivative of *R* with respect to *Z* is

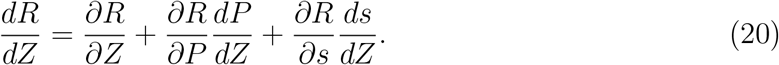

From (19), the first term corresponds to

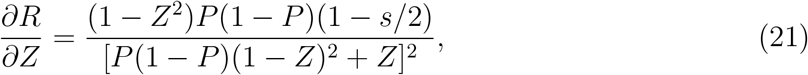

indicating that the partial derivative of *R* with respect to *Z* is negative for *Z >* 1 and positive for *Z <* 1. The partial derivative of *R* with respect to *P* is non-positive:

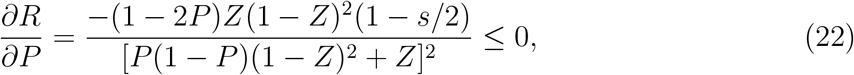

with equality only if

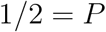

(see (17)). Lastly, *R* declines under higher rates of selfing (*s*):

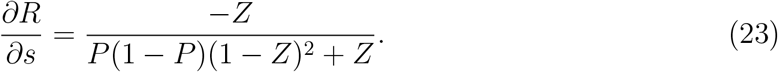

## 3 Dioecious systems of reproduction

A full examination of the nature of the effect of sex-specific viability *Z* on *R* (20) requires determination of the uniparental fraction among reproductives (*s*) and the contribution to the gene pool of gonochores (*P*), as well as an assessment of their dependence on *Z* (*dP/dZ, ds/dZ*). I now obtain explicit expressions for these components for the models of androdioecy and gynodioecy addressed here.

### 3.1 Androdioecy

Consider an androdioecious population comprising *N*_*H*_ hermaphroditic and *N*_*M*_ male reproductives, with males contributing small gametes at rate *σ*_*A*_ relative to hermaphrodites (Figure 1). Under the assumption of a sufficiency of small gametes (sperm or pollen) at any positive frequency of males or hermaphrodites, the uniparental fraction *s*_*A*_ under androdioecy is independent of the proportion of reproductive males. Accordingly, the probability that a random individual of reproductive age is uniparental corresponds to

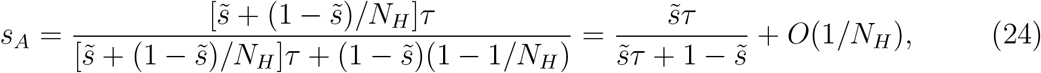

for *τ* the rate of survival of uniparental offspring relative to biparental offspring. To *O*(1*/N*_*H*_), the uniparental fraction under androdioecy and full hermaphroditism (10) are equal. That the uniparental fraction is independent of *Z* implies

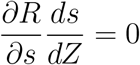

in (20).

From (19), relative effective number at the ESS sex ratio is

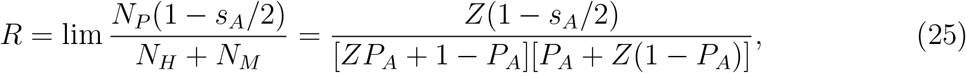

for *Z* the viability of males relative to hermaphrodites and *P*_*A*_ the probability that a gene randomly sampled from an offspring of reproductive age derives from a male parent. Among biparental offspring of reproductive age, the proportion that have a male parent is proportional to *σ*_*A*_:

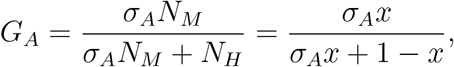

for *x* the proportion of males among reproductives. The reproductive value of a male is

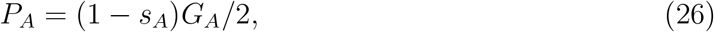

and from a hermaphroditic parent with the complement probability. Expression (26) indicates that among biparental reproductives (1 − *s*_*A*_), a proportion *G*_*A*_ have a male parent, with 1*/*2 the probability that the paternally-derived gene is sampled.

The ESS sex ratio at reproductive age (18) corresponds to

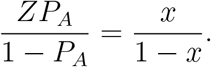

Elimination of *x* from the two expressions for *P*_*A*_ yields the proportion of the gene pool contributed by males at the ESS,

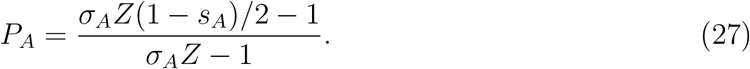

As one might expect, the contribution of males declines with increases in *s*_*A*_, the fraction of offspring that are uniparental. Expression (27) also determines the ESS sex ratio (18) among reproductives:

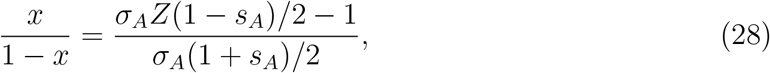

in agreement with Equation (7) of Lloyd (1975). It is valid only if

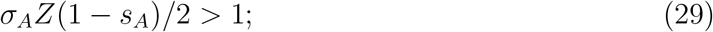

otherwise, the ESS sex ratio corresponds to full hermaphroditism (*x* = 0, Uyenoyama and Takebayashi 2017).

Expression (27) confirms that males contribute less to the gene pool than hermaphrodites,

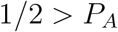

(17), and that *∂R/∂P*_*A*_ (22) is negative. As one might expect, the gonochore contribution to the gene pool increases with gonochore viability

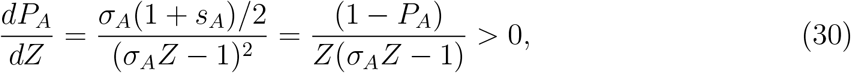

in which *s*_*A*_ is independent of *Z*. We then have

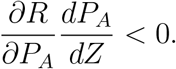

This relation, together with the negativity of *∂R/∂Z* (21) for *Z >* 1, indicates that relative effective number *R* declines as *Z* increases in cases in which males have higher viability than hermaphrodites (*Z >* 1). In this range, *R* declines from its maximum of (1 − *s*_*A*_*/*2), the level expected under full hermaphroditism, as the sex ratio at reproduction departs from the sex ratio at conception in favor of males.

Consider now the remaining case, in which hermaphrodite viability exceeds male viability (1 *> Z*). Because males are maintained only if (29) holds, *Z* lies in the range

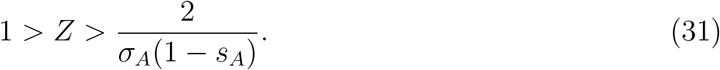

For *Z* near the lower bound, (27) indicates that hermaphrodites contribute nearly the entire gene pool (*P*_*A*_ ≈ 0), implying that *R* approaches its maximum (1 − *s*_*A*_*/*2) at both ends of this range. This behavior suggests non-monotonic dependence of *R* on male viability *Z*. Substitution of (21), (22), and (30) into (20) produces an expression for the derivative of *R* with respect to *Z*. In the range (31), this expression is proportional to a quadratic in *Z*,

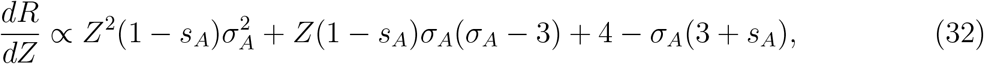

which is positive at *Z* = 1 and negative at *Z* = 2*/*[*σ*_*A*_(1 − *s*_*A*_)], the limits of the range of validity. From the maximum of (1 − *s*_*A*_*/*2) at *Z* = 1, *R* first declines as *Z* declines, reaching its minimum value at the single positive root of this quadratic (32). As *Z* declines further, *R* increases, again approaching its maximum (1 − *s*_*A*_*/*2) as *Z* approaches its lower bound.

### 3.2 Gynodioecy

Now consider a gynodioecious population comprising *N*_*H*_ hermaphroditic and *N*_*F*_ female reproductives (*N* = *N*_*H*_ + *N*_*F*_), in which females produce large gametes at rate *σ*_*G*_ relative to hermaphrodites (Figure 2). As under androdioecy, a proportion 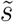 of the large gametes produced by a hermaphrodite are self-fertilized, with the complement fertilized from the small gamete pool of the population. For *τ* the rate of survival of uniparental offspring relative to biparental offspring, the probability that a random individual of reproductive age is uniparental corresponds to

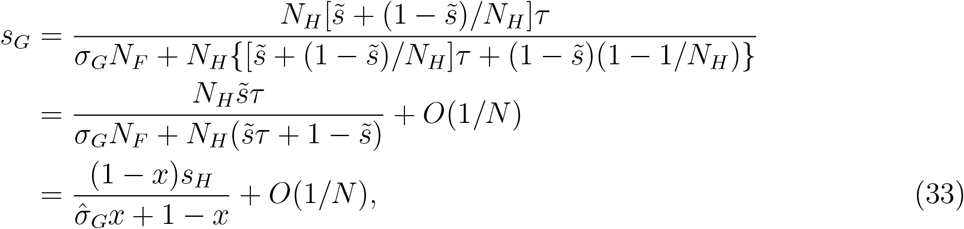

for *s*_*H*_ given in (10), *x* the proportion of females among reproductives, and

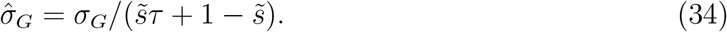

Relative to a hermaphrodite, a female generates 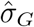 offspring of reproductive age. The presence of obligately-outcrossing females (*x >* 0) reduces the fraction of offspring that are uniparental:

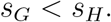

This aspect of gynodioecy contrasts with androdioecy, under which the presence of males has no effect on the uniparental fraction (24).

To large terms, (33) indicates that the proportion of biparental offspring of reproductive age that have a female parent is

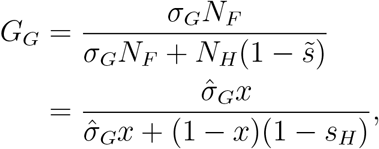

for *x* the proportion of females among reproductives and 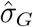 given by (34).

The probability that a gene randomly sampled from the offspring generation at the point of reproduction derives from a female parent (reproductive value of a female) is

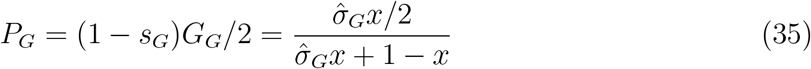

(compare (26)).

The ESS sex ratio at reproductive age (18) is

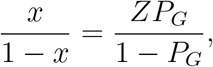

for *x* the proportion of females among reproductives. Substitution of this sex ratio into (35) and solving for *P*_*G*_ produces the proportion of the gene pool contributed by females at the ESS:

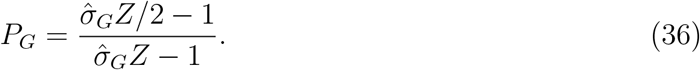

The collective contribution of females to the gene pool increases with the relative viability of females (*Z*), while remaining less than the collective contribution of hermaphrodites (17). The sex ratio among reproductives,

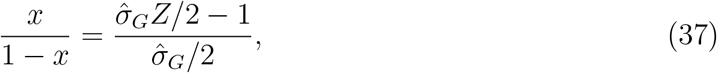

is valid only if

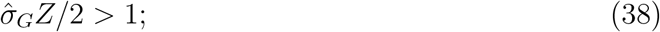

otherwise, the ESS sex ratio corresponds to full hermaphroditism (*x* = 0, Uyenoyama and Takebayashi 2017).

Lloyd (1975) presented a parameter-dense model of gynodioecy, including multiple stages at which self-pollen may fertilize ovules produced by hermaphrodites (“males” in Lloyd’s terminology). Agreement between (37) and Lloyd’s Equation (2) requires that his *o* = 1*/σ*_*G*_, *S* = 1*/Z, i* = *τ*, and *b* = *e* = 1 (equal fertilization rates of ovules produced by females and hermaphrodites). Also required are that either 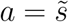 with *w* = 1 or *a* = 0 with 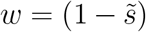, conditions under which the large bracket in Lloyd’s Equation (2) reduces to 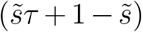, the average viability of zygotes produced through ovules generated by hermaphrodites.

From (19), relative effective number at the ESS sex ratio is

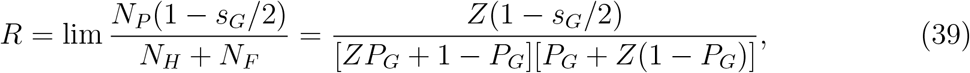

for *P*_*G*_ given by (36). The derivative with respect to *Z* of the collective contribution of females is

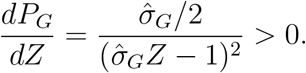

As *∂R/∂P*_*G*_ (22) is negative, the effect of *Z* on relative effective number through the collective contribution of females is also negative:

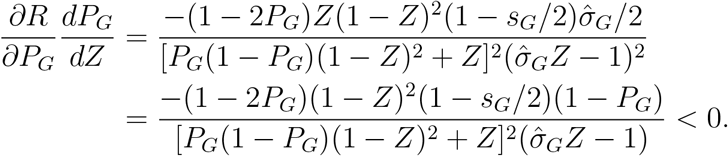

A novel feature of gynodioecy is that the uniparental fraction *s*_*G*_ (33) at the ESS sex ratio (37) depends on *Z*:

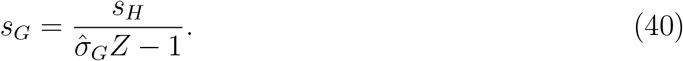

Higher female viability *Z* reduces the uniparental fraction by increasing the proportion of obligately outcrossing reproductives:

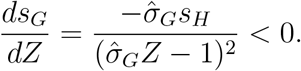

An increase in the number of females tends to promote higher neutral genetic diversity under the ESF (4) through this effect:

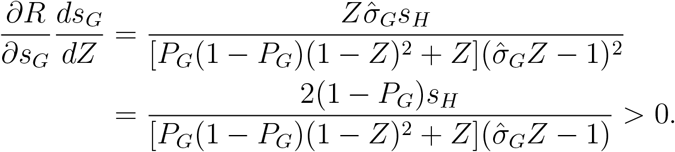

In androdioecious systems, in which the contribution of males is wholly dependent on hermaphrodites (24), higher gonochore viability (*Z*) increases *R* only for *Z* between unity and the positive root of (32). Under gynodioecy, the first terms in (20),

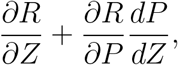

exhibit similar behavior (with *s* = *s*_*G*_). However, the effect of *Z* on *R* through the uniparental fraction,

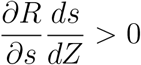

induces qualitatively new aspects into the relationship between *R* and *Z*. While the general finding (19) that relative effective number *R* cannot exceed (1 − *s*_*G*_*/*2) continues to hold, this upper bound increases with *Z*. For *Z* near the minimum value that permits the maintenance of females (38), both *R* and its upper bound (1 − *s*_*G*_*/*2) increase with *Z*. This increase of *R* with *Z* also occurs in the range in which female viability exceeds hermaphrodite viability (*Z >* 1), up to very large values of *Z*.

## 4 Discussion

This analysis addresses the combined effects of a number of factors that influence the level of neutral genetic diversity maintained a population. Among the most important include the level of inbreeding, the intensity of inbreeding depression, the reproductive value of each mating type, differential viability among mating types, and the evolution of the sex ratio. Relative effective number *R* (2) summarizes the overall effect of such factors across the genome. For a given per-generation rate of mutation (*u*) and total number of reproductives (*N*), *R* provides a standard of comparison among full gonochorism, full hermaphroditism, androdioecy, and gynodioecy.

### 4.1 Conservation biology

In an influential article, Franklin (1980) brought the theory of quantitative genetics to bear on conservation biology, reviewing various effective numbers that capture several implications of small population number. To permit the introduction by mutation of new variation underlying quantitative traits to match the pace of its loss genetic drift, Franklin (1980) suggested a minimum effective number of 500, an order of magnitude greater than the conventional guideline at that time based on the direct experience of breeders of domesticated stock. Later characterized as the 50*/*500 rule (Jamieson and Allendorf 2012), these figures appeared to take on a life of their own, sparking much discussion concerning the key criteria relevant to conservation (*e*.*g*., extinction risk, evolvability) and the nature of the genetic data (*e*.*g*., neutral Mendelian markers, components of quantitative variation) that might serve as the basis for the determination of the minimum effective number.

Here, I use *R* (2), a ratio of effective numbers, to compare the capacity of various mating systems to maintain genetic diversity. These effective numbers reflect distinct concepts and in general have distinct expectations (Crow 1954; Crow and Denniston 1988; Ewens 1982). Relative effective number *R* should not be regarded as a substitute for the minimum effective number, be it 500 or larger (Lande 1995). Whether levels of neutral genetic diversity speak to the identification of endangered species, detection of past bottlenecks in population size, or assessment of resilience to major future shifts in habitat or population size (see Teixeira and Huber 2021; Garcia-Dorado and Caballero 2021) lies beyond the objectives of this study. Rather, *R* provides a genome-wide summary of the context in which evolution in a population occurs.

### 4.2 Neutral diversity across reproductive systems

Relative effective number *R* (2) depends on not only the population sex ratio but also the relative viability of mating types (*Z*).

#### Full gonochorism

In the classical case, with reproduction restricted to mating between a pair of mating types in the absence of biparental inbreeding, the effective numbers inferred from rates of parent-sharing and coalescence are identical (*N*_*P*_ = *N*_*C*_) and equal to the harmonic mean of 2*N*_*F*_ and 2*N*_*M*_ (6). Relative effective number (2) corresponds to

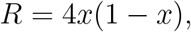

for *x* the proportion of females among reproductives. Irrespective of sex-specific differences in viability (*Z* ≠ 1), the ESS sex ratio at conception is unity (Fisher 1958). Any sex-specific differences in viability (*Z* ≠ 1) that induce departures between the sex ratios at reproduction and at conception diminish relative effective number (*R <* 1).

#### Full hermaphroditism

Self-fertilization reduces the number of parental individuals contributing to the offspring generation. Each uniparental offspring derives from a single parent and each biparental offspring from two distinct parents, implying an average of

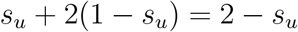

parents per offspring, for *s*_*u*_ the uniparental fraction, the probability that a random reproductive offspring is uniparental. This reduction in the number of reproductives induces correlations between uniting gametes *F* (12) and influences relative effective number:

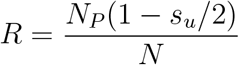

(13). Under full hermaphroditism, the probability that genes sampled from distinct reproductives derive from the same parent is independent of the uniparental fraction (*N*_*P*_ = *N*, Appendix B), implying

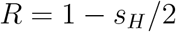

(15), for the uniparental fraction corresponding to *s*_*H*_ (10).

#### Inbreeding under dioecy

Dioecious reproductive systems comprise gonochores together with hermaphrodites. Section 2.4 indicates that relative effective number *R* in general depends on all parameters of the models, including the base rate of selfing 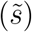, the level of inbreeding depression (*τ*), and sex-specific viability (*Z*). Explicitly,

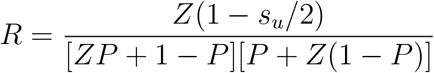

(19) at the ESS sex ratio, for *s*_*u*_ the uniparental fraction and *P* the collective proportion of the offspring gene pool contributed by gonochores.

Under androdioecy (Section 3.1), the uniparental fraction *s*_*A*_ (24) is independent of the frequency of males. This feature reflects the assumption that reproduction at any positive level by hermaphrodites alone provides an abundance of sperm or pollen. Relative effective number *R* takes its maximum value of (1 − *s*_*A*_*/*2) only in the absence of sex-specific viability (*Z* = 1). As *Z* departs from 1 in either direction, *R* tends to decline. However, for *Z* quite low (less than the single positive root of (32)), *R* again approaches its maximum value of (1 − *s*_*A*_*/*2) as the frequency of male reproductives approaches zero.

In gynodioecious populations (Section 3.2), obligate outcrossing by females reduces the uniparental fraction *s*_*G*_ (33) relative to fully hermaphroditic populations (10). By reducing *s*_*G*_, the transition from full hermaphroditism to gynodioecy can increase the level of neutral genetic diversity, behavior that would not be expected in the transition to androdioecy. Unlike any other models studied here, uniparental fraction *s*_*G*_ depends on the relative viability of females (*Z*) as well as the inbreeding depression parameter *τ*. As is the case under androdioecy, relative effective number *R* achieves its maximum value of (1 − *s*_*G*_*/*2) in the absence of sex-specific viability (*Z* = 1). Compared to androdioecy, the dependence of this maximum value on *Z* generates a more complex non-monotone dependence on *Z* across its range.

### 4.3 Empirical estimation of key evolutionary components

Within a model-based context, *R*, a ratio of effective numbers, can be inferred from the pattern of neutral genetic diversity in a random sample (Figure **??**, redelings paper). Redelings paper: ESF used in determining likelihoods based on random samples of microsats from 2 pops of Kmar, gynodioecious plant. Check whether *R* depicted. Uyenoyama and Takebayashi (2017): takes into account ESS sex ratio (check whether *R* or *R/*(1 − *s/*2) shown).

Within a model-based context, relative effective number *R* (2) and other key quantities can be inferred from the pattern of neutral genetic diversity in a random sample. Redelings *et al*. (2015) developed a Bayesian analysis of patterns of variation observed at microsatellite loci in samples derived from a population of the gynodioecious plant *Schiedea salicaria* and two populations of the androdioecious fish *Kryptolebias marmoratus*. Each reproductive system is characterized by a set of basic parameters, including rates of selfing by hermaphrodites 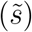, the intensity of inbreeding depression (*τ*), and rates of production of gametes by gonochores (females or males) relative to hermaphrodites (*σ*_*G*_, *σ*_*A*_). The posterior distributions of the basic parameters generated by Redelings *et al*. (2015) permit inference of the posterior distributions of functions of those parameters, including relative effective number (*R*), the collective contribution to the gene pool of gonochores (*P*), and relative viability of gonochores (*Z*).

Figure 3, from Uyenoyama and Takebayashi (2017), depicts the posterior distribution of the proportion of the gene pool contributed by hermaphrodites (1 − *P*), obtained from the analysis of Redelings *et al*. (2015) without imposition of the assumption of convergence of the population sex ratio to the ESS. That practically all the mass of each distribution lies above 0.5 indicates strong support for a greater collective contribution to the gene pool by hermaphrodites compared to gonochores (17). Further, the posterior distributions for *K. marmoratus* suggest that the hermaphroditic contribution is higher in the BP population than in the TC population, concordant with the higher uniparental fraction *s*_*A*_ inferred for the BP population.

**Figure 3:**
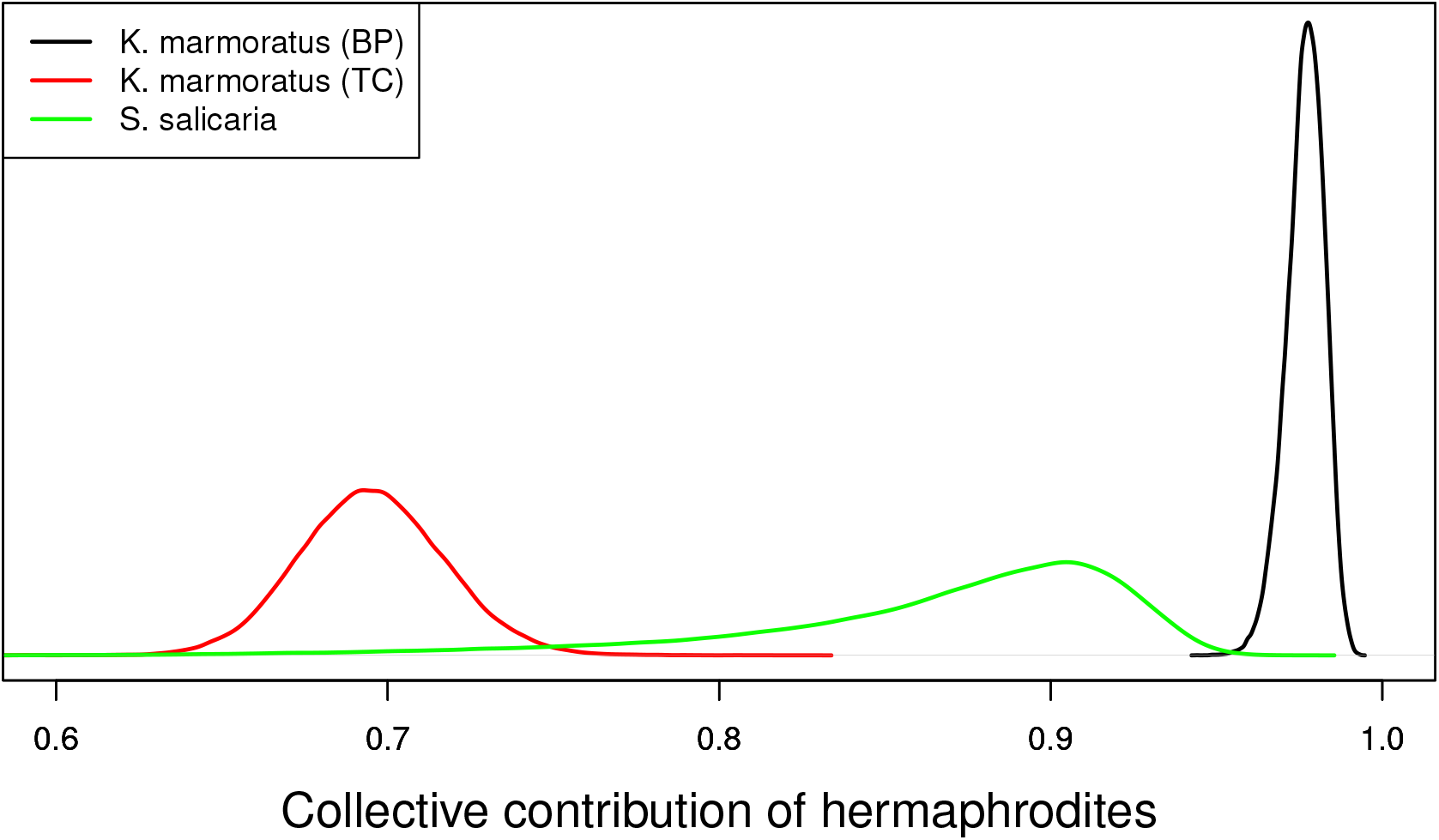
Posterior distributions of collective contribution of hermaphrodites to the gene pool (1 − *P*), inferred from microsatellite data derived from the gynodioecious plant *Schiedea salicaria* and from 2 populations (BP and TC) of the androdioecious fish *Kryptolebias marmoratus* (Redelings *et al*. 2015).

In *K. marmoratus*, an emerging model system for environmental sex determination (ESD, Kelley *et al*. 2016), functional hermaphrodites may transform into fertile males (secondary males). In addition, incubation at lower temperatures promotes the development of fertile primary males directly from self-fertilized eggs (Harrington 1967). Ellison *et al*. (2015) found that interactions between incubation temperature and methylation patterns affect the proportion of primary males, with the source population from which the laboratory lines were derived affecting the nature of the response to experimental treatments. This study appeared to suggest that ESD, mediated by methylation of genes controlling sex expression, may provide a means of regulating the rate of selfing. An alternative view is that the sex ratio evolves in response to the selfing rate. In the model of androdioecy studied here (Figure 1), the ESS sex ratio (28) evolves in response to the uniparental fraction *s*_*A*_ (24), in addition to the relative contribution of males to the sperm pool (*σ*_*A*_) and the viability of males (*Z*). Higher uniparental fractions reduce the collective contribution of males to the offspring generation (26). That quantitative traits, perhaps including the sex ratio under ESD, can evolve rapidly has both conceptual (Sinnott-Armstrong *et al*. 2021) and empirical (Garcia Castillo *et al*. 2024) support.

Central to the present analysis is relative effective number *R* (2), the large-population limit of a ratio of two effective numbers:

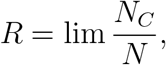

for *N*_*C*_ determined from the rate of coalescence and *N* corresponding to the total number of reproductives. While the estimation of either effective number would require additional information (*e*.*g*., the per-generation rate of mutation *u*), their ratio can be estimated directly (Redelings *et al*. 2015). Figure 11 of (Redelings *et al*. 2015) and Figure 1 of Uyenoyama and Takebayashi (2017) depict *R/*(1 − *s/*2), for *s* the uniparental fraction under androdioecy (24) or gynodioecy (33), without assuming convergence of the population sex ratio to the ESS. (In both papers, the abscissa axis is mislabeled as the relative effective number rather than *R/*(1 − *s/*2).) Imposing the assumption that the *K. marmoratus* populations had in fact evolved to the ESS, Uyenoyama and Takebayashi (2017) generated posterior distributions for *Z*, the viability of males relative to hermaphrodites. Those distributions (Fig. 3 of Uyenoyama and Takebayashi 2017) indicated strong support for a substantial (twofold) reduction in the relative viability of males in both populations, in spite of marked differences in both male frequency and collective contribution of hermaphrodites (Figure 3) between the populations. Turner *et al*. (2006) described *K. marmoratus* males as more conspicuous and reported higher frequencies of males in laboratory-reared populations than in natural populations. Relative to females, males in natural populations may be subject to reduced viability, perhaps as a consequence of predation.

## Acknowledgments

Public Health Service grant GM 37841 provided partial funding for this research.

## Appendix A Labeled coalescent

In the course of proving the Ewens Sampling Formula (Ewens 1972), Karlin and McGregor (1972) described changes in the allele frequency spectrum (3) occurring across generations. Because this description includes the allelic states of genetic lineages, Uyenoyama *et al*. (2019) denoted this argument as the labeled coalescent. Here, we present a summary of the formulation presented by Uyenoyama *et al*. (2019), which addresses changes on the timescale of the most recent evolutionary event rather than generations.

### A.1 Basic recursion

To determine the probability of observing a sample of labeled genes, we consider the relationship between the AFS of a sample of genes (descendant) to the AFS immediately prior to the most recent evolutionary event (ancestor). This most recent evolutionary event may for example correspond to coalescence, mutation, or migration. Let random variable *D* represent the state of the descendant, *A* the state of the ancestor, and *E* the most recent evolutionary event. For a given evolutionary model with parameters **Φ** (5), the likelihood corresponds to

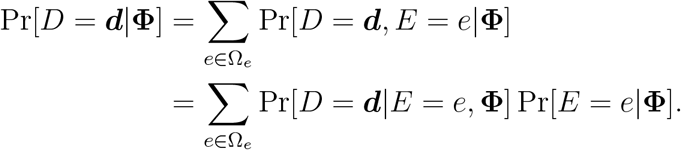

in which ***d*** represents the state of the descendant AFS, *e* a particular realization of the most recent evolutionary event, and Ω_*e*_ the set of all such events. The law of total probability implies

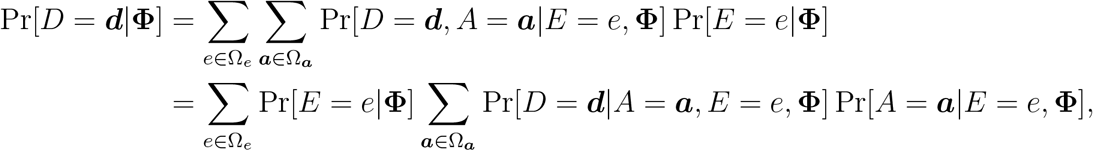

for ***a*** represents the state of the ancestor AFS and Ω_***a***_ the set of all possible ancestor AFSs. Under the assumption that the state of the ancestor *A* is independent of *E*, the evolutionary event forward in time, we have

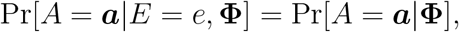

with

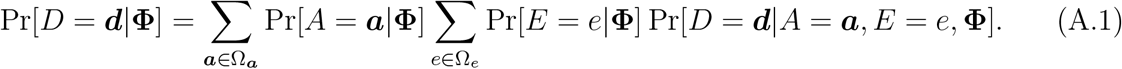

This expression represents a recursion that relates the probability of the present state of the sample (*D*), to the probability of the state immediately prior to the most recent evolutionary event (*A*). In general, determination of the transition term

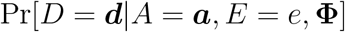

is straightforward: it proceeds from ancestor to descendant through a specified evolutionary event, with only a limited number of ancestor states ***a*** capable of generating descendant state ***d*** through event *e*.

### A.2 Modified ESF

Among independent lineages, the most recent evolutionary event corresponds either to coalescence (*E* = *e*_1_) or to mutation (*E* = *e*_2_), both of which may depend on genomic location or system of reproduction.

For a sample comprising *n* genes, the probabilities that mutation or coalescence correspond to the most recent evolutionary event are

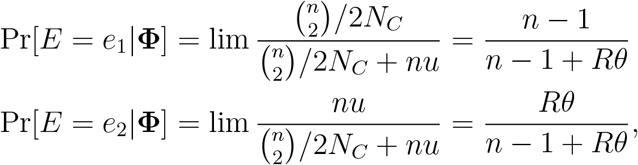

for *θ* (1) and *R* (2) as previously defined.

We first address the case in which coalescence (*E* = *e*_1_) is the most recent event backward in time, which corresponds to the splitting or duplication of a lineage forward in time. The ancestor AFS (*A* = ***a***) comprises only (*n* − 1) genes and has the form

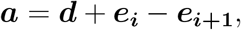

in which ***e***_*j*_ denotes a unit vector, with unity in the *j*^*th*^ position and zeros elsewhere, and *i* is the multiplicity in *A* of the allelic class involved in the splitting event. We define the probability of AFSs with negative elements or other unmeaningful characteristics as zero. A splitting event in an allelic class with multiplicity *i* increases the multiplicity of that allele to (*i* + 1) and decreases by 1 the number of alleles with multiplicity *i*. Using that the split occurs uniformly at random in any of the (*n* − 1) lineages of *A*, we have

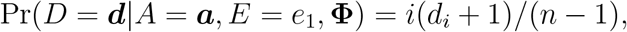

reflecting that (*d*_*i*_ + 1) allelic classes have multiplicity *i* in *A*.

Alternatively, the most recent event may correspond to mutation (*E* = *e*_2_). For an ancestor with AFS identical to the descendant (*D* = *A* = ***d***), the mutation can only have occurred in a lineage representing a singleton allelic class (*i* = 1) in the ancestor AFS, with

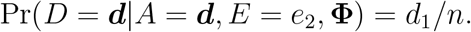

Given this ancestor *A*, a mutation in any other lineage would imply zero probability of generating the descendant *D*. Mutation in an allelic class with multiplicity greater than unity (*i >* 1) generates a new singleton allele and reduces the multiplicity of the allelic class in which the mutation arose. The AFS of ancestors with a positive probability of generating *D* = ***d*** through this route must have the form

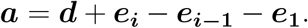

In this case,

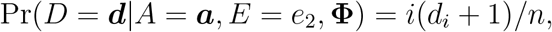

for *i*(*d*_*i*_ + 1) the total number of genes that represent an allele present in multiplicity *i* in the ancestor sample *A*. This expression also reflects that the mutation arises in any gene with equal probability.

At stationarity, the probability of a sample of size *n* with AFS ***a*** is independent of time period:

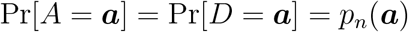

The labeled coalescence recursion (A.1) then reduces to

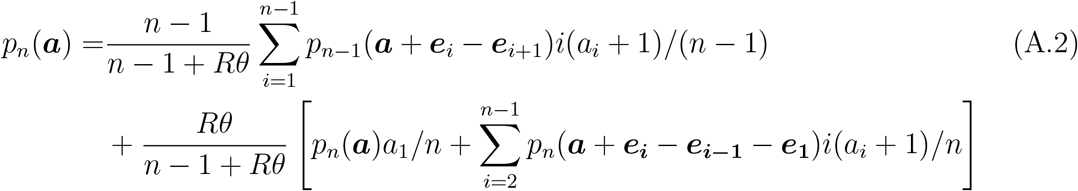

(compare Tavaré 2004), in which unmeaningful expressions (*e*.*g*., probability of spectra with negative elements) are defined as zero.

In the absence of population structure (*R* = 1), Karlin and McGregor (1972) proved the formula given by (Ewens 1972) by showing that it is a solution of (A.2). Some population structures, including those comprising multiple mating types and some forms of regular inbreeding, can be summarized by relative effective number *R* (2). For such cases, the probability of a sample comprising *n* genes with AFS ***a*** corresponds to (4):

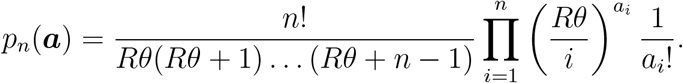

This formula and the probability mass function of the number of alleles observed in a sample of size *n* correspond to Ewens’s (1972) expressions under the substitution of *Rθ* for *θ*. Other remarkable properties of the ESF that are preserved include that the probability that the *n*^*th*^-sampled gene represents a novel allele corresponds to

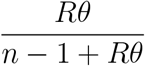

and that the distribution of allele multiplicities conditioned on a given number of alleles in the sample is independent of *Rθ* (Ewens 1972).

Central to the ESF (4) is independence among the sampled lineages. In the presence of inbreeding, lineages of genes sampled from the same organism remain associated by virtue of their shared genealogical history since the most recent outcross. Independence might be ensured by sampling no more than a single lineage per locus per organism. Alternatively, the shared history might be accommodated by regarding sample size (*n*) as a random variable. At the most recent point at which the ancestral lineages of all sampled genes reside in distinct organisms, *n* in (4) corresponds to the number of lineages remaining after coalescence between lineages during the generations of inbreeding until the most recent outcross.

## Appendix B Parent-sharing under full hermaphroditism

This section addresses the per-generation rate of parent-sharing (1*/N*_*P*_) in a population of *N* reproductive hermaphrodites of which a proportion *s*_*H*_ (10) are uniparental. The rate of parent-sharing corresponds to the probability that a pair of genes, each randomly sampled from distinct individuals in the offspring generation, derive from the same parent.

A pair of uniparental offspring share their parent with probability

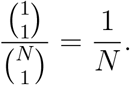

A uniparental offspring and a biparental offspring share a parent with probability

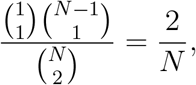

corresponding to the probability that 1 of the parents of the biparental individual is identical to the parent of the uniparental individual. A pair of biparental offspring may have 2, 1, or 0 parents in common. The probability that the second individual shares both parents with the first individual is of order 1*/N* ^2^,

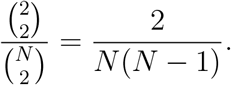

It shares exactly one parent with probability

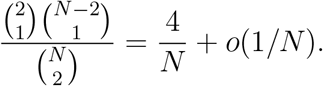

To order 1*/N*, the probability that a random pair of offspring share exactly one parent corresponds to

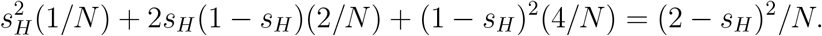

Given parent-sharing between the pair of offspring individuals, the probability that 2 genes, 1 sampled from each individual, derive from the shared parent is 1 for two uniparental individuals, 1*/*4 for two biparental individuals, and 1*/*2 in the remaining case. In agreement with (14), the probability that a pair of genes, each sampled from distinct reproductives in the offspring generation, derive from the same parent is

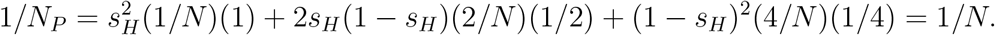

Under full hermaphroditism, this index of effective number is independent of the uniparental fraction *s*_*H*_.

